# Does the seed fall far from the tree? Weak fine-scale genetic structure in a continuous Scots pine population

**DOI:** 10.1101/2023.06.16.545344

**Authors:** Alina K. Niskanen, Sonja T. Kujala, Katri Kärkkäinen, Outi Savolainen, Tanja Pyhäjärvi

## Abstract

Knowledge of fine-scale spatial genetic structure, *i.e.*, the distribution of genetic diversity at short distances, is important in evolutionary research and in practical applications such as conservation and breeding programs. In trees, related individuals often grow close to each other due to limited seed and/or pollen dispersal. The extent of seed dispersal also limits the speed at which a tree species can spread to new areas.

We studied the fine-scale spatial genetic structure of Scots pine (*Pinus sylvestris*) in two naturally regenerated sites located 20 km from each other in continuous south-eastern Finnish forest. We genotyped almost 500 adult trees for 150k SNPs using a custom made Affymetrix array. We detected some pairwise relatedness at short distances, but the average relatedness was low and decreased with increasing distance, as expected.

Despite the clustering of related individuals, the sampling sites were not differentiated (*F_ST_* = 0.0005). According to our results, Scots pine has a large neighborhood size (*Nb* = 1680– 3210), but a relatively short gene dispersal distance (*σ*_g_ = 36.5–71.3 m). Knowledge of Scots pine fine-scale spatial genetic structure can be used to define suitable sampling distances for evolutionary studies and practical applications. Detailed empirical estimates of dispersal are necessary both in studying post-glacial recolonization and predicting the response of forest trees to climate change.

## Introduction

Understanding the fine-scale spatial genetic structure of species is important in a wide range of fields. The structure builds up from the interaction of gene flow and extrinsic factors, such as fragmented habitats and differences in growth conditions, which in turn affect selection, population size and density. On the other hand, intrinsic factors, such as dispersal ability and mating patterns of the species also have an influence (Loveless & Hamrick 1984; Vekemans & Hardy 2004). Gene flow takes place between individuals in physical space (Bradburd & Ralph 2019), often leading to an isolation-by-distance pattern (Wright 1943; Málecot 1967). Thus, individuals close to each other are commonly expected to be more closely related than a random sample from within the species. In sedentary species, such as trees, spatial aggregation of related individuals often results from limited seed and/or pollen dispersal (Hardy & Vekemans 1999). In wind-pollinated species, pollen can disperse long distances, from hundreds of meters to hundreds of kilometers (e.g., Kremer *et al*. 2012; Desilva & Dodd 2021), whereas seed dispersal has shorter average distances (Kremer *et al*. 2012). Pollen dispersal distances in animal-pollinated species are commonly shorter—from a few meters to a few kilometers (e.g., Levin & Kerster 1974; Kremer *et al*. 2012; but see Ahmed *et al*. 2009). As a result of the differences in dispersal distances, strong fine-scale spatial genetic structure is common in animal-pollinated and rare in wind-pollinated tree species (Vekemans & Hardy 2004; Hardy *et al*. 2006; Born *et al*. 2008; Vakkari *et al*. 2020). The distribution of genotypes at short distances is usually transient, largely impacted by life-history traits and can be evaluated through estimating relatedness between individuals.

In evolutionary research, spatial genetic information is valuable for inferring the strength of the evolutionary forces—genetic drift, selection, and gene flow—that participate in forming the genetic structure (Rousset 2003; Slatkin 1985). The fine-scale spatial genetic structure of populations needs to be considered in many applications too. For example, in genotype-phenotype association analyses, spurious associations may arise due to correlation of allele and trait frequencies in different populations, but also within a population, if the underlying genetic structure is not corrected for (Pritchard & Rosenberg 1999; Persyn *et al*. 2018). In practical applications, where individuals are chosen from natural populations for conservation, population management or breeding programs, it is essential to know how genetic diversity and, for example, rare alleles are distributed in space to maintain high genetic diversity, avoid inbreeding and, on the other hand, unintended mixing of differentially adapted populations (Desilva & Dodd 2021; Escudero *et al*. 2003; Smith *et al*. 2018). When the span of spatial autocorrelation is known, sampling can be adapted to the needs of each application.

Knowledge of fine-scale spatial genetic structure can recursively be used to infer dispersal distances (Málecot 1967; Rousset 1997 & 2003). The ability to disperse becomes increasingly important in the light of climate change, as plant populations may become maladapted to their current locations (Gougherty *et al*. 2021). Dispersal information can be used, for example, in predicting species’ potential for adaptation (Kuparinen *et al*. 2010; Kremer *et al*. 2012; Barton 1979; Slatkin 1973), bearing in mind that dispersal rates in, e.g., the open landscapes of colonization stage may be different than those estimated here. If the natural dispersal rate is estimated to be too slow for responding to climate change, human assisted migration is one possible way to aid adaptation (Marris 2009; Aitken & Whitlock 2013).

Scots pine (*Pinus sylvestris*) is a keystone conifer species in large parts of the forests of Northern Eurasia and, therefore, important for ecosystem functioning (Pyhäjärvi *et al*. 2020 and references therein). As a major source of timber, paper and pulp, Scots pine also has high economic value, especially in Fennoscandia (e.g., in Finland, https://www.luke.fi/en/statistics/wood-consumption/forest-industries-wood-consumption-2021). Due to its continuous and wide distribution, wind-pollination and predominantly outcrossing mating system (Muona & Harju 1989), Scots pine shows very weak genetic population structure at the global scale, largely in the form of subtle isolation-by-distance across its distribution range (Tyrmi *et al*. 2020). Less is known about within-population genetic structure as only small, fragmented populations have been studied in this respect so far (Robledo-Arnuncio & Gil 2005; Sofletea *et al*. 2020). Although artificial regeneration with genetically improved seedlings has become more common in forestry, natural regeneration has been the predominant regeneration method in forests, making the patterns described here common across Fennoscandia. Since Scots pine has such a dominant role in the boreal forest ecosystems, even small changes in its distribution or adaptive ability may have large consequences.

Here we investigate the fine-scale spatial genetic structure of Scots pine in two naturally regenerated sites in south-eastern Finland. We use a genome-wide 400k single nucleotide polymorphism (SNP) array (Kastally & Niskanen *et al*. 2022), which allows us to estimate relatedness in a large sample of 469 trees. We estimate the parameters of the isolation-by-distance model and use this information to derive estimates of gene dispersal distance. We then investigate the spatial spread and sharing of rare alleles that can have different fitness effects and average allele age compared to more common alleles. The knowledge of fine-scale spatial genetic structure and dispersal distance is useful in practical applications of Scots pine breeding and in associating genetic and trait variation in natural populations, but also for modeling adaptation and making predictions on how widely distributed wind-pollinated trees can respond to climate change.

## Material and Methods

### Samples and genotypes

Our study population at the Punkaharju intensive study site (ISS, https://www.evoltree.eu/resources/intensive-study-sites/sites/site/punkaharju) in south-eastern Finland includes two sampling sites, Mäkrä (61°50’16.8’’N, 29°23’39.5’’E) and Ranta-Halola (61°39’19.7’’N, 29°17’14.7’’E) located 21 km apart (Figure 1). The landscape in south-eastern Finland has mostly continuous pine forest cover, lakes and some agricultural areas. The study site stands arose through natural regeneration after seed tree cuttings (i.e., retaining only part of the mature trees to provide seed for establishing the next generation) 60–70 years ago in Mäkrä and an unknown, but probably a few decades longer, time ago in Ranta-Halola. We sampled 469 adult Scots pines at approximately 20 m distances, with the shortest within-sampling site distance between trees being 10 and 14 m and the longest 464 and 1164 m in Mäkrä and Ranta-Halola, respectively. We selected trees that were similar in size by eye—113 trees from Mäkrä and 356 trees from Ranta-Halola. The mean age, estimated by counting the tree rings from a core sample at breast height, was 60.6 (range: 33–112.5) and 90.3 (range: 43–144.5) years in Mäkrä and Ranta-Halola, respectively (Figure S1). We could not estimate the age for two trees from Ranta-Halola.

**Figure 1.**
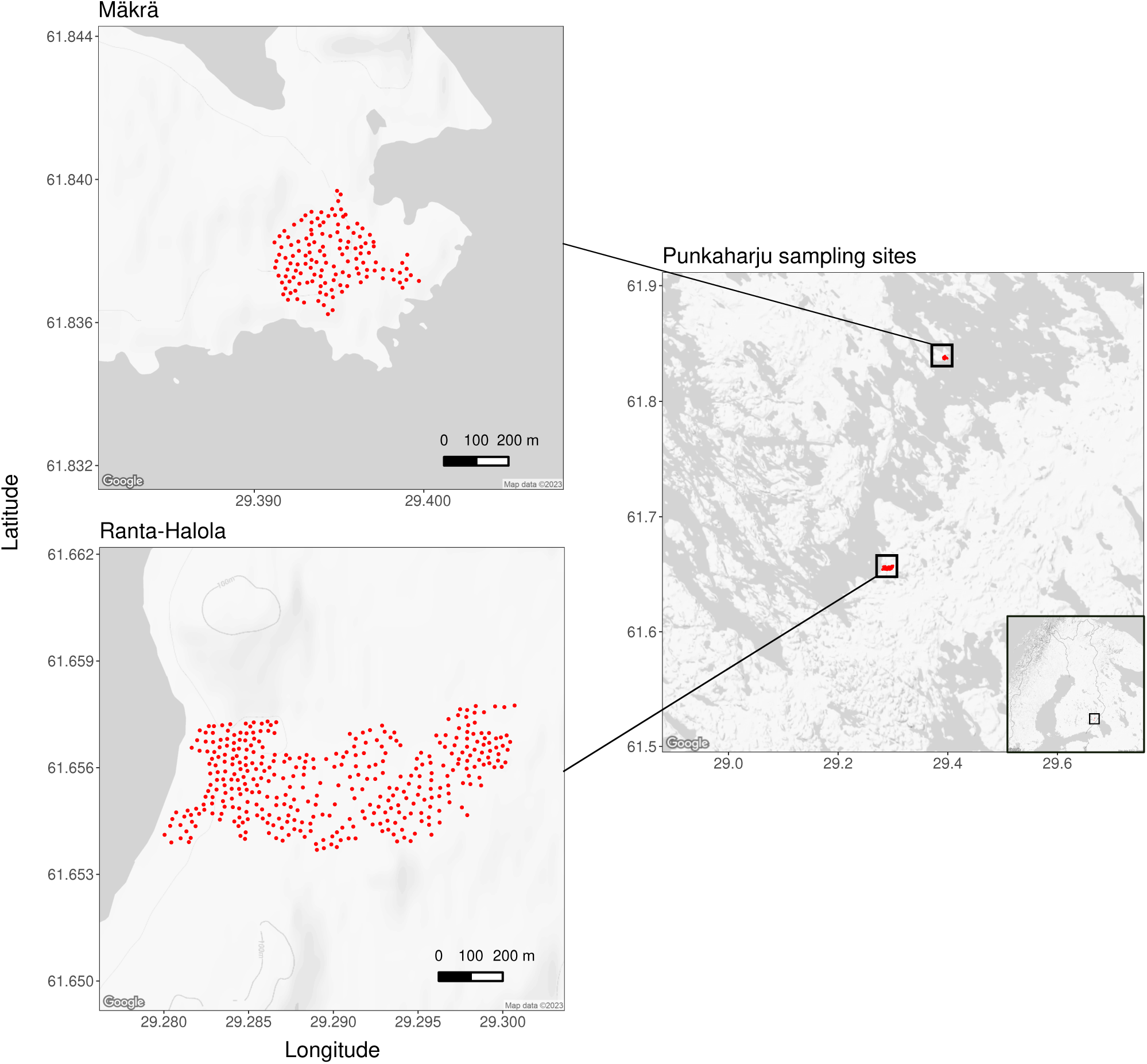
Maps of the sampling sites in the Punkaharju intensive study site located in south-eastern Finland. Sampled trees are indicated as red dots. Mäkrä and Ranta-Halola are located 21 km apart. The maps were drawn in R using the package *ggmap* (Kahle & Wickham 2013).

Needle samples from the adult trees were genotyped on a custom-made Affymetrix SNP array including 407 540 SNPs. Development of the SNP array and genotyping of the samples is described in detail in Kastally & Niskanen *et al*. (2022). In short, of the 407 540 SNPs we used a dataset of 157 325 polymorphic SNPs with the ThermoFisher conversion types Poly High Resolution (three well-separated genotype clusters) and No Minor Homozygote (two well-separated genotype clusters, homozygous and heterozygous) as a starting point for filtering the loci. We used four embryo samples and their parents (described in Kastally & Niskanen *et al*. 2022) to estimate Mendelian errors for each locus using PLINK (v. 1.9; Purcell *et al*. 2007) and excluded SNPs with more than one Mendelian error. Since pines have a high proportion of repetitive and paralogous genome sequence (Wegrzyn *et al*. 2014), we excluded SNPs with more than one seemingly heterozygous genotype in haploid megagametophyte samples (described in Kastally & Niskanen *et al*. 2022) to avoid SNPs in potentially paralogous genomic regions. Further filtering of the genotype data was done according to the requirements of each analysis as described below.

### Spatial data and pairwise distances

We recorded the coordinates for each tree using a portable GPS locator in 2020. The initial coordinates in the ETRS-TM35FIN geodetic coordinate system were transformed to the coordinates in the EUREF-FIN-GRS80 geodetic coordinate system using a Finnish map service on the web (https://kartta.paikkatietoikkuna.fi/). One of the study trees from Ranta-Halola had died between the needle sample collection and coordinate recording and was excluded from the spatial analyses. We estimated the pairwise spatial distance matrix for 468 adult trees with coordinates using the R (v. 3.6.3, R core team 2020) package *fields* function “rdist.earth” (Nychka *et al*. 2017).

### Population structure

To get an overall picture of the genetic structure of our study population, we conducted principal component analysis (PCA) using the R package *pcadapt* (Privé *et al*. 2020). We ran PCA for two sets of individuals, first for all 469 individuals, and second for 332 individuals excluding: i) individuals used in the SNP discovery (Kastally & Niskanen *et al*. 2022), ii) individuals related to the SNP discovery individuals (pairwise relatedness, genomic relationship matrix (GRM) ≥ 0.044; Yang *et al*. 2011; see details below), and iii) one individual from each pair with pairwise relatedness ≥ 0.044. We excluded closely related individuals and SNP discovery individuals from the PCA to detect the underlying population structure without the signal of family structure or SNP ascertainment effects. We used a set of 65 498 SNPs with the following characteristics: minor allele frequency (MAF) ≥ 0.05, close to Hardy Weinberg equilibrium (HW; exact test *p* value ≥ 0.001), and not in high linkage disequilibrium (LD) r^2^ < 0.9 with other SNPs in 10 kb windows within a contig (using PLINK), unless stated otherwise.

We estimated pairwise *F_ST_* (Weir & Cockerham 1984) between the two study sites using the R package S*tAMPP* (Pembleton *et al*. 2013) and performed 1000 bootstraps to estimate its 95% confidence interval (CI).

### Fine-scale spatial genetic structure

We estimated pairwise relatedness between all 469 samples as a genomic relationship matrix (GRM) using the GCTA (Yang *et al*. 2011) command “--make-grm-gz”. GRM was estimated between individuals *j* and *k* over SNP loci from *i* to *N* using the formula

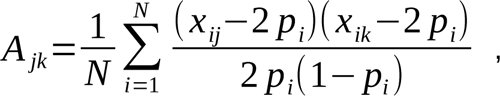

where *xij* and *xik* are the numbers of reference (major) alleles in individuals *j* and *k*, and *p_i_* is the reference allele frequency. Since inclusion of closely related individuals may inflate the relatedness estimates, we used allele frequencies estimated for 387 unrelated adult individuals (GRM < 0.0625, the mean of relationship class for, e.g., first cousins once removed) as reference allele frequencies in the estimation of GRM. We classified each pair of individuals into family relationship classes (e.g., see Manichaikul *et al*. 2010 for kinship relationship estimates (*F*), which are equal to half of the corresponding relatedness estimates) based on pairwise GRM: second degree between [0.177,0.354), e.g., half-siblings, third degree between [0.088, 0.177), e.g., first cousins, fourth degree between [0.044, 0.088), e.g., first cousins once removed, and unrelated below 0.044. We also estimated the genomic inbreeding coefficient (*F_GRM_*) for each individual based on genomic relationship matrix in GCTA.

The Mantel test (Mantel 1967) is a traditional test of spatial autocorrelation where the relationship of two dissimilarity (i.e., distance) matrices is investigated. In spatial genetics, the null hypothesis of a Mantel test is that genetic distance, or similarity when measured as relatedness or kinship, and spatial distance are not correlated. To study the relationship between relatedness and spatial distance, we estimated their correlation using Spearman’s correlation coefficient (*ρ*) separately for the two sampling sites Mäkrä and Ranta-Halola. We conducted the Mantel test using the R package *ecodist* (Goslee & Urban 2007) with 10 000 permutations. The use of the Mantel test in spatial genetics has been criticized because of the requirements of homoscedasticity and linear correlation between genetic and spatial distances for the data (Legendre *et al*. 2015). These problems are, however, less severe in the Mantel correlogram analysis where samples are divided into pre-defined spatial distance classes and each distance class is compared separately to joint data from other distance classes. We therefore also used Mantel correlograms (10 000 permutations) to evaluate the correlation between relatedness and distance within distance classes.

### Neighborhood size and dispersal distance

We estimated neighborhood size (*Nb*), the effective number of potentially mating individuals belonging to a within-population neighborhood (Wright 1946), and gene dispersal distance (*σ_g_*) using an iterative approach implemented in SPAGeDi (Hardy & Vekemans 2002). To obtain *σ_g_*, SPAGeDi first estimates a starting value for neighborhood size using the formula

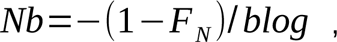

where *F_N_* is the mean kinship (Loiselle *et al*. 1995) of the first distance class (0–50 m in Mäkrä and 0–60 m in Ranta-Halola) and *blog* is the slope of the regression of kinship on the natural logarithm of spatial distance over all distance classes (Rousset 2000; Hardy & Vekemans 2002). Kinship was estimated with 28 378 SNPs with MAF ≥ 0.20 (due to computational limitations), using the formula

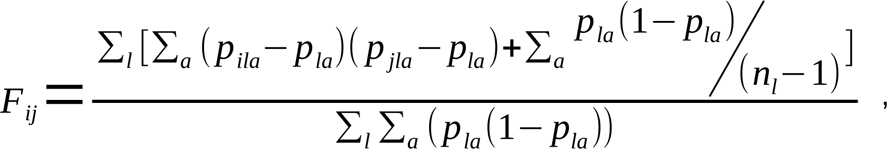

where *p_ila_* and *p_jla_* are the frequencies of allele *a* at locus *l* in individuals *i* and *j*, *p_la_* is the reference allele frequency of allele *a* at locus *l*, and *n_l_* is the number of gene copies defined in the sample at locus *l* (Loiselle *et al*. 1995; Hardy & Vekemans 2002). Then gene dispersal distance was estimated using the formula

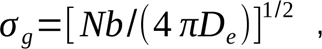

where *D_e_* is the effective population density that accounts for differences in the reproductive success of individuals (Hardy & Vekemans 2002). The *Nb* estimation procedure was repeated using *blog* from kinship∼distance regression up to the distance of the chosen maximum σ value. Census density (*D*) estimates of unmanaged Finnish Scots pine forests vary between 608–4470 trees per hectare (Lönnroth 1926), and the census density in commercial forests is about 2 000 trees/ha (200 000 trees per km^2^; Fahlvik *et al*. 2005; Väisänen *et al*. 1989). Our study sites have regenerated naturally after cutting to an unknown, but likely lower than 2000 trees/ha density of seed trees. However, as pollen and seed dispersal from surrounding forests (Jiménez-Ramírez *et al*. 2021) likely increases the effective density, we used 2000 trees/ha as our starting point, and estimated *σ_g_* assuming ratios of effective to census density of 0.25 (*D_e_* = 500 trees/ha) and 0.5 (*D_e_* = 1 000 trees/ha). Gene dispersal distance can reliably be estimated within a distance that is assumed to be in mutation-drift-equilibrium that should be reached in a few generations within the distance *σ_g_*/(2*µ*)^1/2^ (*µ* = mutation rate; Rousset 1997 & 2000). Based on a 10^-3^ mutation rate of microsatellites, this distance is approximately 20\**σ_g_*. Since the SNP mutation rate (10^-9^; Willyard *et al*. 2007) is lower than the microsatellite mutation rate, we estimated *Nb* (and thus also sigma) using both 20\**σ_g_* and a higher value of 60\**σ_g_*. However, since the maximum distances within our study sites are shorter than the estimated 20\**σ_g_* distance, increasing the maximum distance does not affect our estimates of sigma nor *Nb*. To measure the strength of the fine-scale spatial genetic structure, we estimated the intensity of spatial genetic structure (*Sp*; Vekemans & Hardy 2004) using the formula

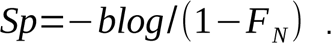

We used 1 000 permutations of individual locations in estimating *blog* to test how probable it is to get higher *Sp* by chance.

### Rare alleles

To study the spatial spread of rare alleles, we investigated sharing of rare alleles with MAF < 0.01 (23 623 SNPs after removing singletons) between individuals and correlated (Spearman’s *ρ*) this with pairwise relatedness and spatial distance. To avoid the SNP discovery ascertainment bias in rare allele sharing, we estimated the correlation using only 426 individuals excluding: i) individuals used in the SNP discovery and ii) individuals with pairwise relatedness (GRM) >= 0.044 with the SNP discovery individuals. We used within sampling site Mantel correlograms to study the correlation between rare allele sharing and spatial distance within distance classes. *P*-values for the correlations were obtained with Mantel’s test using 10 000 permutations. For illustrative purposes, we fitted a local (LOESS) regression in R for the proportion of shared rare alleles on relatedness and included sampling site as a fixed predictor.

## Results

### Spatial genetic analyses

On the population scale with all 469 adult trees, we identified a very weak structure between Mäkrä and Ranta-Halola based on PCA (Figure 2). The majority of the samples in both Mäkrä and Ranta-Halola clustered together along principal components (PC) 1 and 2 (Figure 2a). However, we detected one outlying cluster of samples from Ranta-Halola on PC1 and one on PC2, and a group of samples from Mäkrä dispersed along PC2. These outlying samples consisted of related individuals, and the outliers disappeared when we conducted PCA only for 332 individuals that were unrelated (GRM < 0.044) and not used in the SNP discovery (Figure 2b). Concordantly, we found a low F_ST_ of 0.0005 (95% CI: 0.0004–0.0005) between Mäkrä and Ranta-halola.

**Figure 2.**
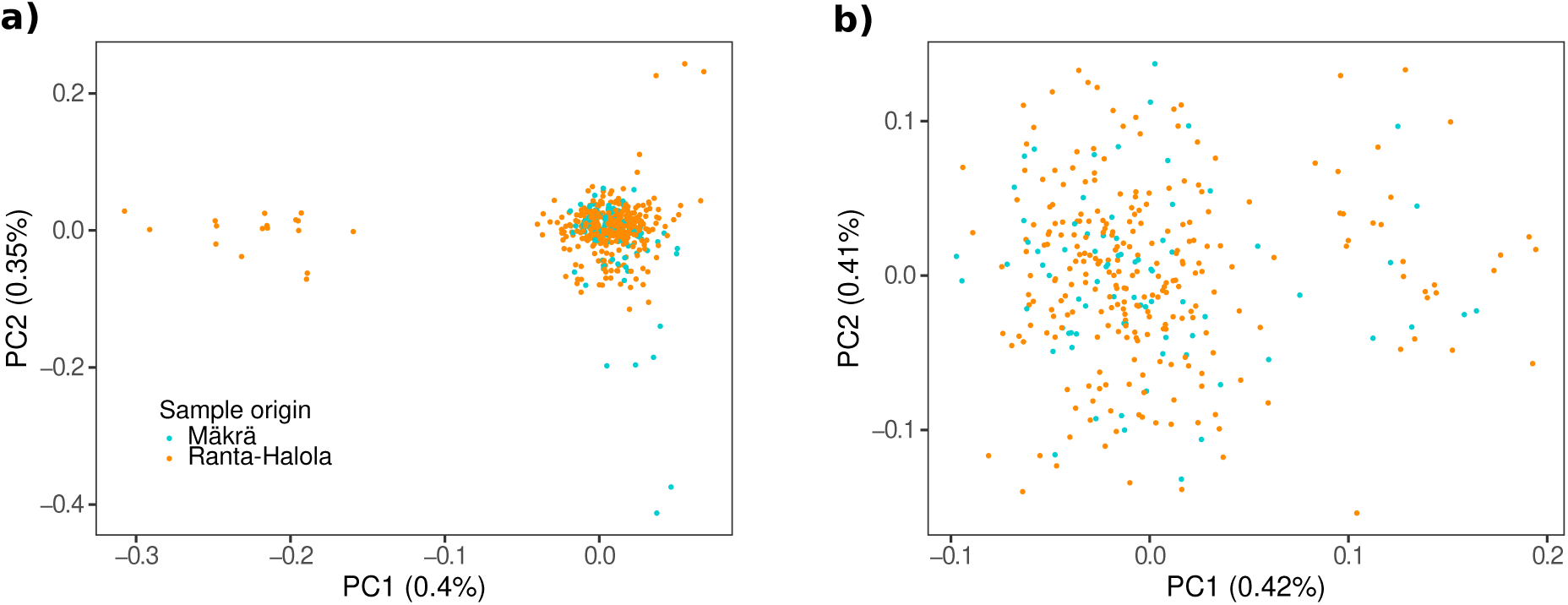
Principal component analysis (PCA) for Mäkrä and Ranta–Halola for a) all 469 adult trees and b) only 332 trees; excluding individuals used in the SNP discovery (Kastally & Niskanen *et al*. 2022), individuals with pairwise relatedness higher than 0.044 with the SNP discovery individuals, and one individual of each pair with pairwise relatedness ≥ 0.044.

We found that pairwise relatedness was low within the study sites (Figure 3; Table S1). Even in the class with the shortest distance between individuals, the mean GRM was only 0.004 in Mäkrä and 0.002 in Ranta-Halola. However, relatedness still decreased with increasing distance in both sampling sites at a similar rate (Figure 3; see Figure S2 for distance restricted to Mäkrä’s maximum distance). We used relatedness (GRM) or kinship (*F*, Loiselle) to estimate pairwise relatedness depending on the analysis, and both methods gave similar estimates of relatedness (Pearson’s *r* = 0.842, p < 0.0001; Figure S3). The spatial genetic structure was evident in closer distance classes until ∼65 m in Mäkrä and ∼200 m in Ranta-Halola. In these distance classes, there was a negative correlation between relatedness and distance as shown by the Mantel correlograms (Figure 4; Figure S4). The Mantel test indicated a subtle but significant decay of relatedness with spatial distance at Ranta-Halola (Spearmann’s *ρ* = -0.044, one-tailed *p* < 0.0001) but not at Mäkrä (Spearmann’s *ρ* = -0.010, one-tailed *p* = 0.216). We also estimated the intensity of spatial genetic structure and found that, even when there was evidence for spatial genetic structure, its intensity was low in both study sites (Ranta-Halola *Sp* = 0.0008, one-tailed *p* < 0.001, and Mäkrä *Sp* = 0.0005, one-tailed *p* < 0.001). The low intensity was caused by the small decrease in the pairwise kinship with distance over each sampling site and by the low average kinship in the first distance class.

**Figure 3.**
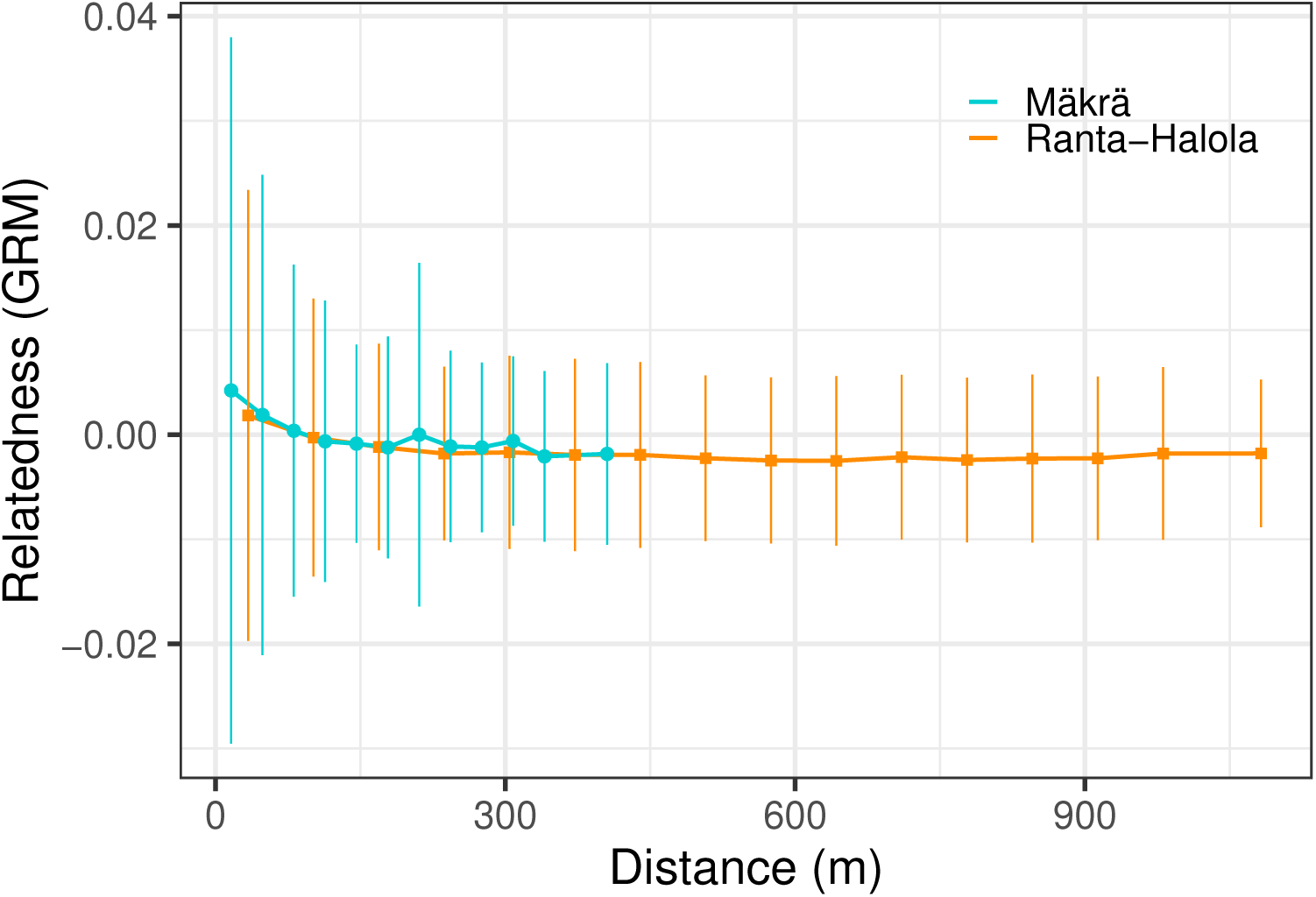
Decay of pairwise relatedness with distance in Mäkrä (turquoise) and Ranta-Halola (orange) sampling sites. Mean (circles and squares) and standard deviation (vertical lines) of relatedness is plotted for each distance class; the number of pairwise comparisons, the mean GRM and the mean distance in each distance class are shown in Table S1.

**Figure 4.**
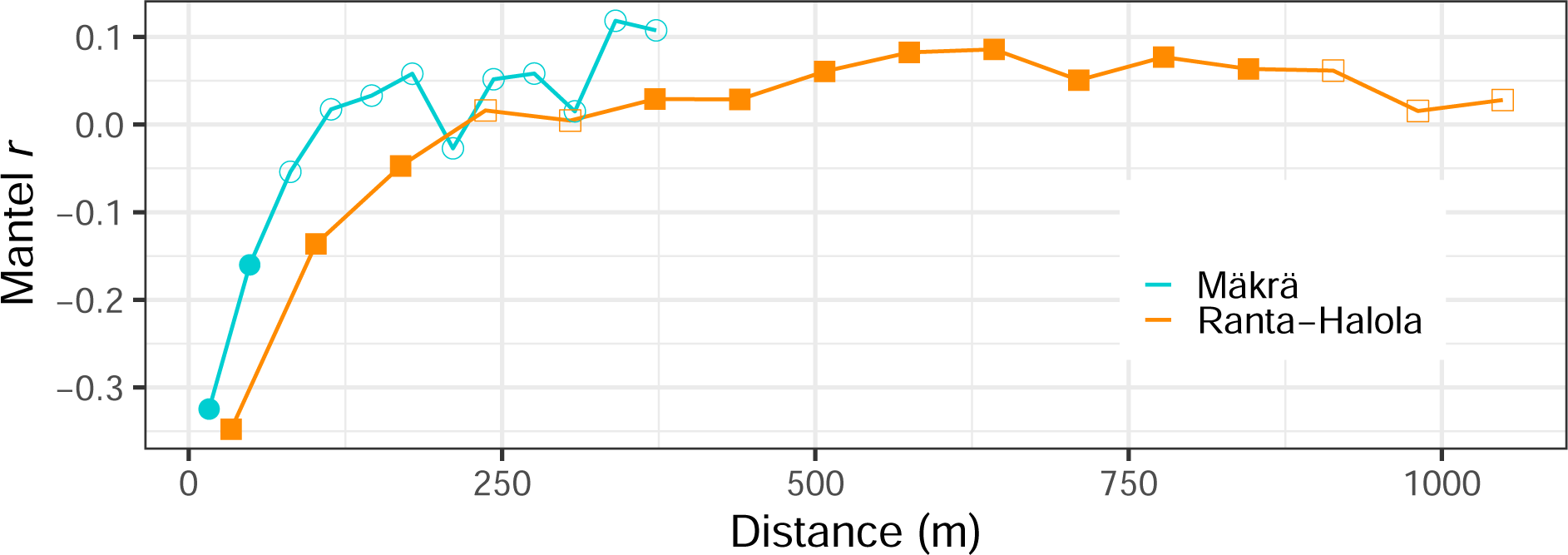
Mantel correlogram showing the correlation (measured as Mantel *r*) between pairwise relatedness and distance within each distance class in a) Mäkrä (circles, turquoise) and b) Ranta-Halola (squares, orange). Filled shape indicates a *p*-value < 0.05. Two of the longest distance classes from Mäkrä and one from Ranta-Halola have been left out due to including less than 100 pairwise comparisons.

Among the 69 163 pairwise comparisons, 24 closely related pairs (GRM ≥ 0.177) were identified (Figure 5). All related individuals (GRM ≥ 0.044) were growing in the same sampling site. The highest relatedness (GRM) for a between sampling site pair was 0.031, whereas the highest within site relatedness was 0.332 in Mäkrä and 0.349 in Ranta-Halola. When the trees were divided into family relationship classes, the median distance of the most related family relationship class found here (GRM = 0.177–0.354, indicating second-degree relatedness) was 51 m in Mäkrä and 59 m in Ranta-Halola, compared to the respective median distances of 166 m and 357 m for unrelated individuals (GRM < 0.044; Figure 5). This illustrates that the spatial aggregation of closely related individuals is similar in both the smaller and the larger study area. Family relationship classes categorized using relatedness vs. kinship estimates were very similar, with a few pairs categorized in the neighboring classes (Table S2). We did not find any sign of close inbreeding in our sample; the highest inbreeding coefficient (*F_GRM_*) was 0.035 in Mäkrä and 0.080 in Ranta-Halola.

**Figure 5.**
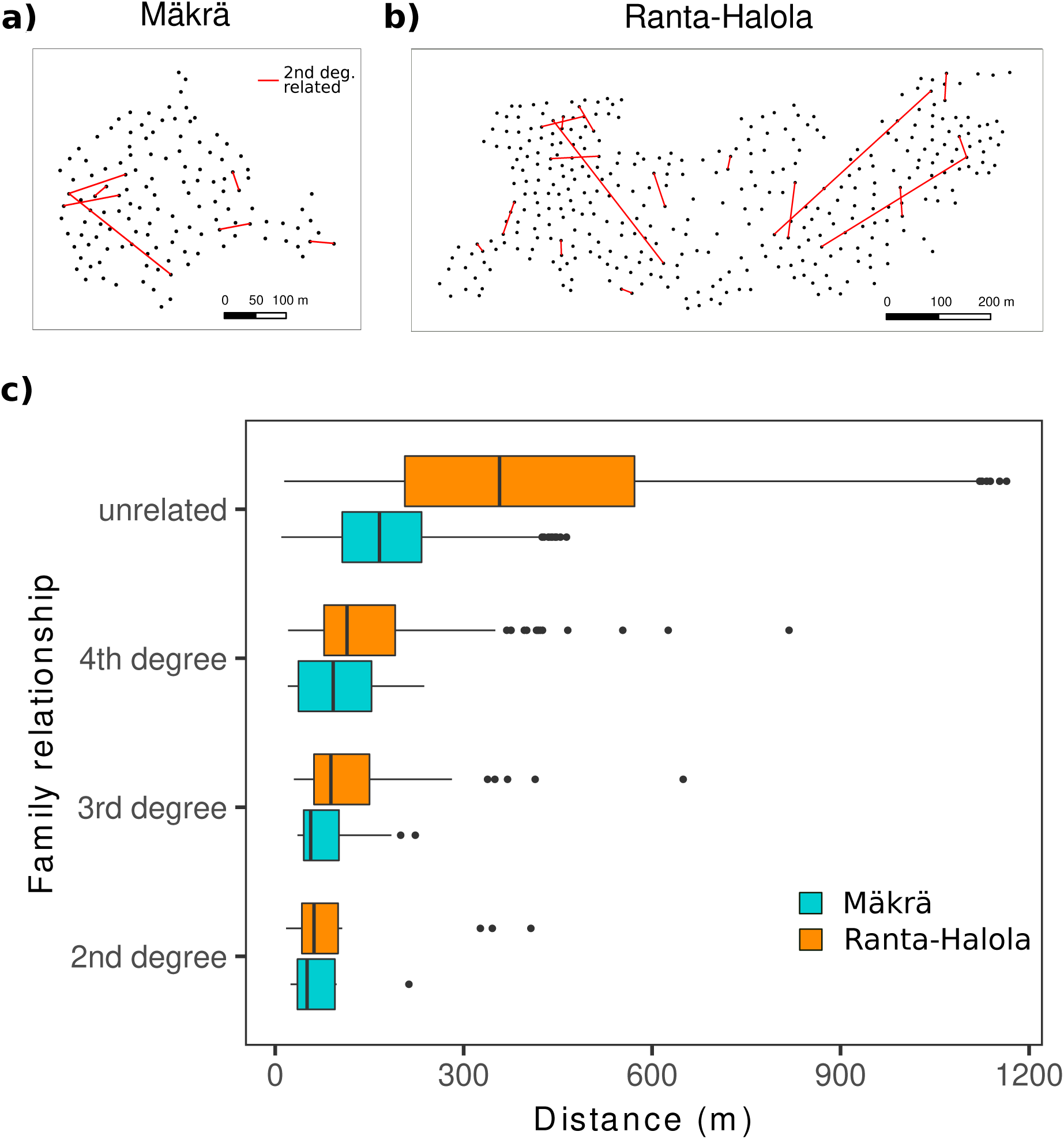
Association between family relationship classes and spatial distance. Pairwise distances of the second degree related individuals in a) Mäkrä and b) Ranta-Halola are shown as red lines. The distribution of all pairwise distances of individuals in different family relationship classes are shown in c) for Mäkrä (turquoise) and Ranta-Halola (orange). Boxplots show the median (central vertical line), the lower and upper quantiles (boxes), and up to 1.5 interquartile range (whiskers) distances. Family relationships are classified based on pairwise GRM: second degree between 0.177–0.354 (e.g., half-sibling; n = 7 in Mäkrä and n = 17 in Ranta-Halola), third degree between 0.088–0.177 (e.g., first cousin; n = 12 in Mäkrä and n = 38 in Ranta-Halola), fourth degree between 0.044–0.088 (e.g., first cousin once removed; n = 33 in Mäkrä and n = 164 in Ranta-Halola), and unrelated below 0.044 (n = 6276 in Mäkrä and n = 62 616 in Ranta-Halola).

Depending on the effective population density (*D_e_*) estimate we used, the mean neighborhood size (*Nb*) over the two Scots pine sites was 3210 (with *D_e_* = 500 trees/ha; Table 1) or 1680 trees (with *D_e_* = 1 000 trees/ha). We estimated the mean gene dispersal distances (σ_g_) jointly with *Nb* and found that the mean σ_g_ were 71.3 m (*D_e_* = 500 trees/ha; Table 1) and 36.5 m (*D_e_* = 1 000 trees/ha).

**Table 1.** Gene dispersal distance (*σ_g_*) and neighborhood size (*Nb*) estimates for Mäkrä and Ranta-Halola for two different ratios of effective (*D_e_*) to census (*D*) density. In this study, *D_e_/D* ratio of 0.5 equals *D_e_* = 1 000 trees/ha and 0.25 equals *D_e_* = 500 trees/ha. Examples of *Nb* and *σ_g_* for animal- and wind-pollinated tree species from previous studies below.

| Study site<br>(*Pinus sylvestris*) | $D_e / D = 0.5$ | | $D_e / D = 0.25$ | | Pollinator | Reference |
| --- | --- | --- | --- | --- | --- | --- |
| | $N_b$ | $\sigma_g$ (m) | $N_b$ | $\sigma_g$ (m) | | |
| Mäkrä | 1404 | 33 | 3673 | 76 | wind | this study |
| Ranta-Halola | 1957 | 39 | 2747 | 66 | wind | this study |
| **Species** |  |  |  |  |  |  |
| *Dicorynia guianensis* | 116 | 222 |  | 203 | insects | Hardy *et al.* 2006 |
| *Moronobea coccinea* | 20 | 134 |  | 195 | birds | Hardy *et al.* 2006 |
| *Milicia excelsa* |  |  | 370 | 3755 | wind | Bizoux *et al.* 2009 |
| *Pinus pinaster* * | 97 | 30 |  |  | wind | De-Lucas *et al.* 2009 |
| *Araucaria angustifolia* | 381 | 140 |  |  | wind | Sant'Anna *et al.* 2013 |
| *Thuja occidentalis* * | 65 | 55 |  |  | wind | Pandey & Rajora 2012 |

\*Only core or continuous populations included.

### Rare alleles

In line with overall relatedness, we found that more related individuals also shared a higher proportion of rare alleles than unrelated individuals (n = 426*, ρ* = 0.049, one-tailed *p* < 0.0001 estimated using Mantel’s test; Figure 6), but this relationship was only visible in higher relatedness values. Rare allele sharing and distance had a weak negative relationship within study sites (*ρ* = -0.010, *p* = 0.057 in Ranta–Halola and *ρ* = -0.028, *p* = 0.084 in Mäkrä; Figure S5). Mantel correlograms showed that pairwise sharing of rare alleles decreased with increasing spatial distance—similarly to the decrease of relatedness with spatial distance (Figure S6).

**Figure 6.**
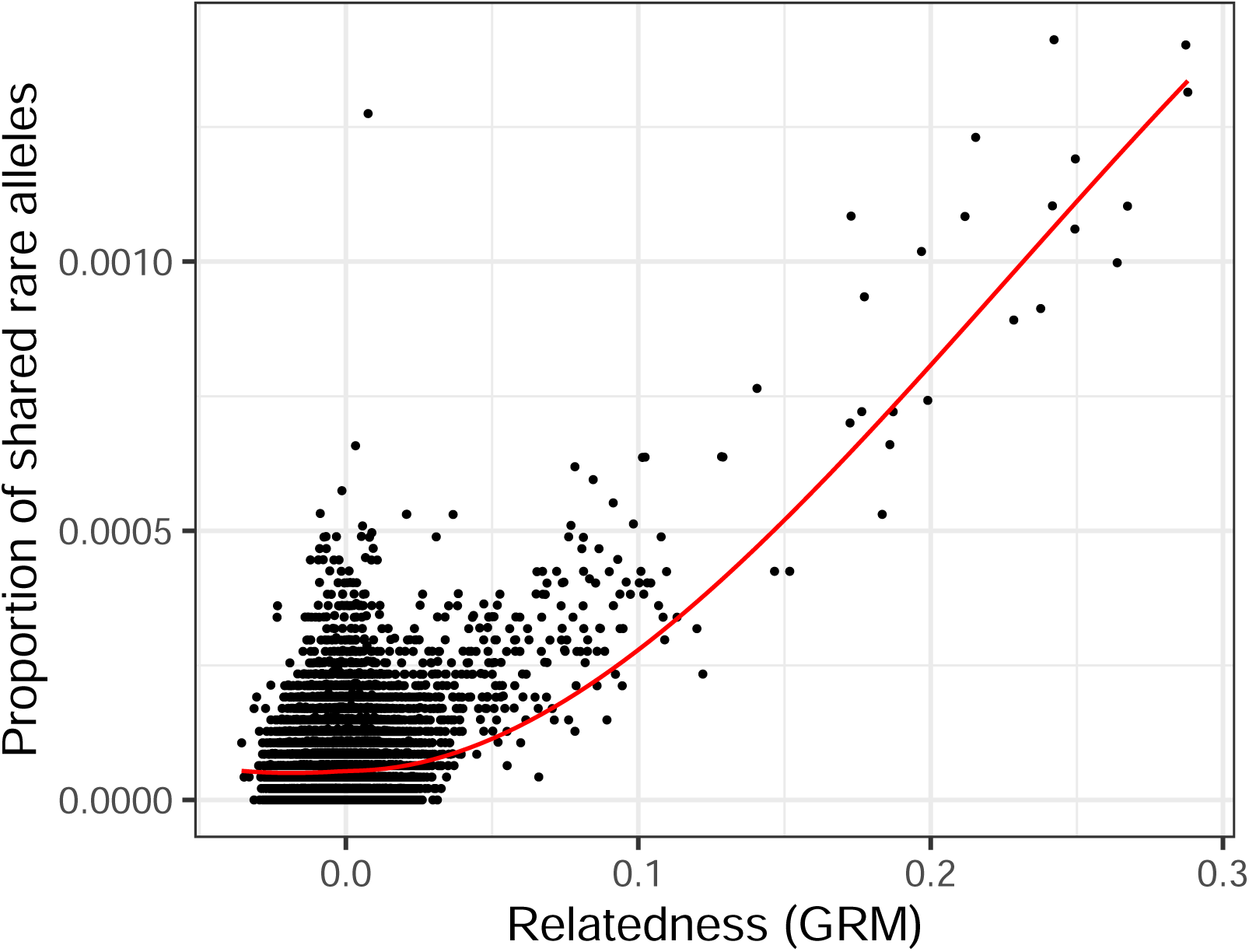
Relationship between proportion of shared rare alleles of all rare alleles and relatedness (GRM). Pairwise comparisons between individuals (dots) and LOESS curve fitted to the proportion of shared rare alleles on relatedness (GRM).

## Discussion

### Large continuous Finnish population of Scots pine has weak fine-scale genetic structure

Species with a very subtle structure among populations are often considered panmictic, i.e., mating completely randomly also within populations. While Scots pine genotypic frequencies follow HW frequencies at the adult stage similarly to a panmictic species (Muona & Harju 1989, Pyhäjärvi *et al*. 2020), the mating patterns are not completely random in space (Robledo-Arnuncio & Gil 2005; Torimaru *et al*. 2012). Concordant with spatially restricted mating patterns, we showed that fine-scale spatial genetic structure— albeit very weak—is maintained in adult Scots pine stands (Figures 3 & 4). This was evident in the spatial proximity of individuals with higher pairwise relatedness (Figure 5). Scots pine has partial selfing (5–10% of mature seeds), but only a part of the selfed offspring survive to the mature seed stage (Koski 1971, Kärkkäinen & Savolainen 1993), and even fewer survive to the adult stage (Koelewijn *et al*. 1999). The average mortality of the selfed seeds is 75–85%, compared to 20–30% for the open-pollinated seeds (Kärkkäinen *et al*. 1996, Koelewijn *et al*. 1999). In our samples, we did not find high inbreeding estimates or pairwise relatedness values close to 1, which are expected in selfed progeny. Thus, relatedness patterns of adult trees found here are not a result of selfing. Fine scale genetic structure has previously been found in smaller fragmented Scots pine populations (Robledo-Arnuncio & Gil 2005; Sofletea *et al*. 2020), but here we show this pattern for the first time within a large continuous population.

We found that the spatial genetic structure reached somewhat longer distance in Ranta-Halola than Mäkrä (Figure 4; Figure S4). This may partly be caused by differences in the shape and size of the sampled areas. In addition, each distance class includes considerably more pairwise comparisons in Ranta-Halola, which results in more statistical power. The mean age of Ranta-Halola is higher (Figure S1), which allows multiple generations of dispersal events after the seed tree cuttings were done and, therefore, a longer extent of spatial genetic structure. On the other hand, Mäkrä had proportionally more pairs in the second degree family relationship class, which could be caused by the more recent or possibly more intense seed tree cutting. There are likely differences in the fecundity of individual trees (Torimaru *et al*. 2012), which can cause differences between the sites. Nevertheless, the patterns of relatedness (Figures 3 & 5) and spatial genetic structure were very similar between the study sites, which suggests that the dispersal patterns are similar between the sites.

Long-distance pollination events and the continuous distribution, allowing constant gene flow, are likely major contributors to the extremely low intensity of spatial genetic structure we detected (*Sp* from 0.0005 to 0.0008). In addition, the high population density increases the number of potential breeders, and all of these factors keep the fine-scale spatial genetic structure weak even when the average gene dispersal distance is relatively short. In contrast, when populations are far from each other and disjunct, the chance of pollination from nearby trees is higher, which leads to stronger spatial genetic structure. This effect of population fragmentation is evident in previous Scots pine studies where *Sp* has ranged from up to 0.0098 in Scotland (González-Díaz *et al*. 2017) to as high as 0.0207 in the Carpathian Mountains (Sofletea et al. 2020). Our study sites showed low intensity of spatial genetic structure also when compared to other conifers (*Sp* = 0.001–0.0349; Desilva & Dodd 2021; Kitamura *et al*. 2018; Sant’Anna *et al*. 2013; De-Lucas *et al*. 2009; Vekemans & Hardy 2004) and other tree species (*Sp* = 0.002–0.075; Bizoux *et al*. 2009; Hardy *et al*. 2006; Vekemans & Hardy 2004). The low intensity of spatial genetic structure is in line with strong gene flow through pollen. It also indicates that Scots pine populations even at the northern distribution edge receive ample gene flow, and have high genetic variation as long as they are connected to the main population. Varis *et al*. (2009) showed that the Finnish Scots pine populations at different latitudes are receptive for pollen before their own pollen shedding starts, but the pollen from more southern populations is already available for fertilization. This potential for gene flow from southern to northern populations also facilitates adaptation to climate change. However, the situation is different in populations that are more fragmented and isolated, such as the Spanish and Scottish populations, where the pollen dispersal is more restricted and, thus, the spatial genetic structure is stronger (Hampe & Petit 2005).

### Relatively short dispersal distance of Scots pine and its implications for selection

We estimated that the average gene dispersal distance in our Scots pine stands was 53.9 meters (average of 71.3 m and 36.5 m), which is relatively short compared to the gene dispersal distance estimates of other wind-pollinated trees, typically between 30 and 3755 meters (Table 1; Pandey & Rajora 2012; Sant’Anna *et al*. 2013; Bizoux *et al*. 2009; De-Lucas *et al*. 2009), and of animal-pollinated trees up to 1296 meters (Hardy *et al*. 2006). However, population density (estimate) plays a large role in estimating the dispersal distances. While estimating population density in Scots pine stands is straightforward, it is more complicated to assess the density of a forest in the past, i.e., during establishment of seedlings that resulted in the current stand. As an early succession species, regeneration mainly occurs in pulses after disturbances (e.g., forest fires, storms and loggings; Linder *et al*. 1997; Lundqvist *et al*. 2017). Studies on regeneration of seed and shelterwood stands also suggest that seedling establishment of Scots pine occurs in sparse stands (50–200 trees/ha; Beland *et al*. 2010; Rautio *et al*. 2023). In the estimation of gene dispersal distances, the effective density estimate should also take into account the breeding contribution of adult trees surrounding the cut area. It should also be noted that the effective density of the study sites has varied greatly over time, and trees germinating 33 or 143 years ago have faced very different fertilization and germination conditions. We used the effective densities of 500 and 1000 trees/ha. By assuming lower effective density, the estimated dispersal distance would increase. However, a very long average dispersal distance does not seem probable given the relatively sharp decay of the mean relatedness with distance and the observed pairwise distances of the related individuals (Figures 3, 4 & 5). Further, earlier Scots pine seed (10–20 m; Debain *et al*. 2007) and pollen dispersal (47.6–53 m; Koski 1970, based on radioactively labelled pollen; Robledo-Arnuncio & Gil 2005) estimates are relatively close to our dispersal distance estimate, bearing in mind that our estimate includes both seed and pollen components. Most dispersal distance estimates—including ours—assume that the distribution of dispersal distances (i.e., the dispersal kernel) is Gaussian. However, especially pollen is able to disperse very long distances (Lindgren *et al*. 1995; Robledo-Arnuncio 2011) leading to potentially more leptokurtic dispersal kernels that are difficult to estimate. These would result in higher mean dispersal distances (Robledo-Arnuncio & Gil 2005; Debain *et al*. 2007). Taken together, our gene dispersal estimates can be taken as minimum estimates given the potential for lower effective density and leptokurtic pollen dispersal.

Knowing the dispersal distance of a species is crucial for predicting how quickly it can spread to new habitats but also for estimating its ability to respond to selection and to adapt. The balance between the amount of gene flow and the strength of selection defines the probability of local adaptation as a response to spatially diversifying selection, as gene flow from differentiated populations causes migration load and hinders local adaptation (gene swamping; Lenormand 2002). Selection can be very efficient in species with large *N_e_*, such as Scots pine, and phenotypic climatic adaptation to different latitudes is well known (Mikola 1982; Aho 1994; Notivol *et al*. 2007; Kujala & Knürr *et al*. 2017). Early mortality is very high, and even when it is largely random, it also provides much opportunity for selection. However, the question is not just how strong the selection is but also on what spatial scale the species can track the environmental differences and adapt through changes in allele frequencies. According to Slatkin (1973), a population can respond to selection only if the underlying environmental heterogeneity occurs over a distance longer than the characteristic length (*L* = *σ / √s*, where *σ* is the offspring mean dispersal distance and *s* is the strength of selection). With our estimates of mean gene dispersal distance (53.9 m) and hypothetical selection coefficients *s* = 0.01 or *s* = 0.001, the characteristic length would be 539 m and 1 705 m, respectively.

No evidence for fine-scale adaptation has been found in Scots pine. Reciprocal transplant experiments showed no evidence of local adaptation to different soils at the scale of some kilometers (Jimenez-Ramírez *et al*. 2023). Furthermore, a nine-year common garden study with progeny of Punkaharju ISS showed that selection at the local population scale was rather weak on the adaptive seedling traits, even if fitness was lower in populations from further north and south (Kujala *et al*. 2023). As adaptive traits are often polygenic, selection on individual loci would be expected to be rather weak, yielding little potential to respond to different selection in close-by sites. In some other European and Mediterranean conifers, considerable selection coefficients were reported for individual loci across steep ecological gradients, with at least 1 km distance and often hundreds of meters of altitudinal difference (Scotti *et al*. 2023).

### Fitness and practical implications of the fine-scale spatial genetic structure

Aggregation of relatives leads to a higher probability of inbreeding and inbreeding depression. Scots pine carries a high number of lethal equivalents (Koski 1971; Savolainen *et al*. 1992), which makes selfing and more distant forms of inbreeding detrimental. Our results indicate that related individuals carry an excess of shared rare alleles. A large number of loci in Scots pine have very low minor allele frequencies (Tyrmi *et al*. 2020). Rare alleles are typically young and also enriched for recessive deleterious variants. Thus, spatial genetic structure may lead to more homozygosity and fitness reduction than expected in a totally panmictic population, where these alleles would seldom appear as homozygotes.

Fine-scale spatial genetic structure also has practical implications, e.g., in tree breeding. When closely located trees are also more likely to be related, determining a suitable collection distance of potential breeding individuals is very important in order to avoid introduction of related individuals into breeding programs. Possible inbreeding also needs to be avoided in production and deployment populations. Due to strong inbreeding depression, manifesting as lowered yield of viable seeds and reduced viability and growth of the seedlings, accidental selection of related individuals to seed orchards would cause problems. Furthermore, information on spatial genetic structure can help to define a minimum distance between trees to be used for collecting seed or cuttings in practical gene conservation work. Knowledge on the extent and intensity of fine-scale spatial genetic structure is of importance also for forest management of naturally regenerating sites as it can guide the optimization of distance between spared seed trees during harvesting.

### Conclusions

Here, we described in detail the extent of fine-scale spatial genetic structure and average dispersal distance in a large population of Scots pine from a continuous part of the distribution. We demonstrated that even a wind-pollinated widely distributed species with large effective population size can have detectable, although weak, fine-scale spatial genetic structure. Our estimates of dispersal distance are relevant for practical applications, predicting responses to environmental changes, and understanding the balance between gene flow and other evolutionary factors, especially selection.

## Acknowledgements

This project was funded by: the European Union’s Horizon 2020 research and innovation program, under grant agreement no. 773383 (to UOULU and Luke); the Seventh framework program for research and development, under grant agreement no. 211868 (to UOULU and Luke); the EU Network of Excellence EVOLTREE grant no. 016322 (to UOULU and Luke); the Academy of Finland grants 287431, 293819 and 319313 (to TP), 307582 (to OS), 307581 (to KK), and 309978 (to STK); NoE EVOLTREE (to STK). We thank Natural Resources Institute Finland staff in Loppi and Punkaharju for collecting samples and recording the GPS positions of the trees, Soile Alatalo for molecular laboratory work at the Ecology and Genetics Unit of the University of Oulu, and Coen Westerduin for comments on the previous version of the manuscript. We are grateful for the PCI Evol Biol recommender Myriam Heuertz and the three reviewers Jean-Baptiste Ledoux, Roberta Loh and Joachim Mergeay for their comments that improved the paper substantially.

## Supplementary Data

The following supplementary data is available as a separate document:

**Figure S1.** The distribution of Scots pine age in Mäkrä (turquoise; *n* = 113) and Ranta-Halola (orange; *n* = 354) sampling sites.

**Figure S2.** Decay of pairwise relatedness with distance in Mäkrä (turquoise) and Ranta-Halola (orange) sampling sites. Mean (circles and squares) and standard deviation (vertical lines) of relatedness is plotted for each distance class; number of pairwise comparisons in each distance class are shown in Table S1. The estimates of Ranta-Halola have been moved two metres forward in the plot to avoid overlap with the estimates of Mäkrä.

**Figure S3.** Pairwise relatedness (GRM; Yang *et al*. 2011) plotted against pairwise kinship (Loiselle; Loiselle et al. 1995) estimated for 468 Scots pines in the Punkaharju research area. The red line shows the expected relationship of 2:1 for these estimates.

**Figure S4.** Correlation (measured as Mantel *r*) between pairwise relatedness and distance within each distance class estimated as Mantel correlogram in a) Mäkrä (circles, turquoise) and b) Ranta-Halola (squares, orange). Filled shape indicates a *p*-value smaller than 0.05. Ranta-Halola and Mäkrä are divided into equally long distance classes. Two of the longest distance classes from Mäkrä and three from Ranta-Halola have been left out due to including less than 100 pairwise comparisons.

**Figure S5.** Decay of the proportion of shared rare alleles with spatial distance in Mäkrä (turquoise) and Ranta-Halola (orange) sampling sites. Mean (circles and squares) and standard deviation (vertical lines) of relatedness is plotted for each distance class.

**Figure S6.** Mantel correlogram for rare allele sharing and pairwise distance in a) Mäkrä and b) Ranta-Halola. Filled circles indicate *p*-value smaller than 0.05.

**Table S1.** The mean and standard deviation (SD) of pairwise relatedness (GRM) in each distance class for Ranta-Halola and Mäkrä sampling sites (Figure 3; Figure S2). The last two and three distance classes for Ranta-Halola (upper panel) and Mäkrä, respectively, are combined so that each class has at least 100 comparisons. The lower panel for Ranta-Halola shows the relatedness values for Ranta-Halola, when distances are classified according to Mäkrä’s 14 distance classes.

**Table S2.** Comparison of family relationship classes estimated for 468 Scots pines in the Punkaharju research area using relatedness (GRM; Yang *et al*. 2011) or kinship estimate (Loiselle; Loiselle et al. 1995). Estimates on the darker green background show the same degree of family relationship and ligher green shows one degree difference in the estimated class. Relatedness was estimated using 65 498 SNPs with MAF ≥ 0.05 and kinship using 28 378 SNPs with MAF ≥ 0.20 due to computational reasons.

## Data availability statement

The data supporting the findings of this study are available in Figshare (DOI: 10.6084/m9.figshare.23531142; the data will be open to public after this manuscript has been accepted for publication).

