## Supplementary Material for "Does the seed fall far from the tree? Weak fine-scale genetic structure in a continuous Scots pine population"


### Supplementary Figures

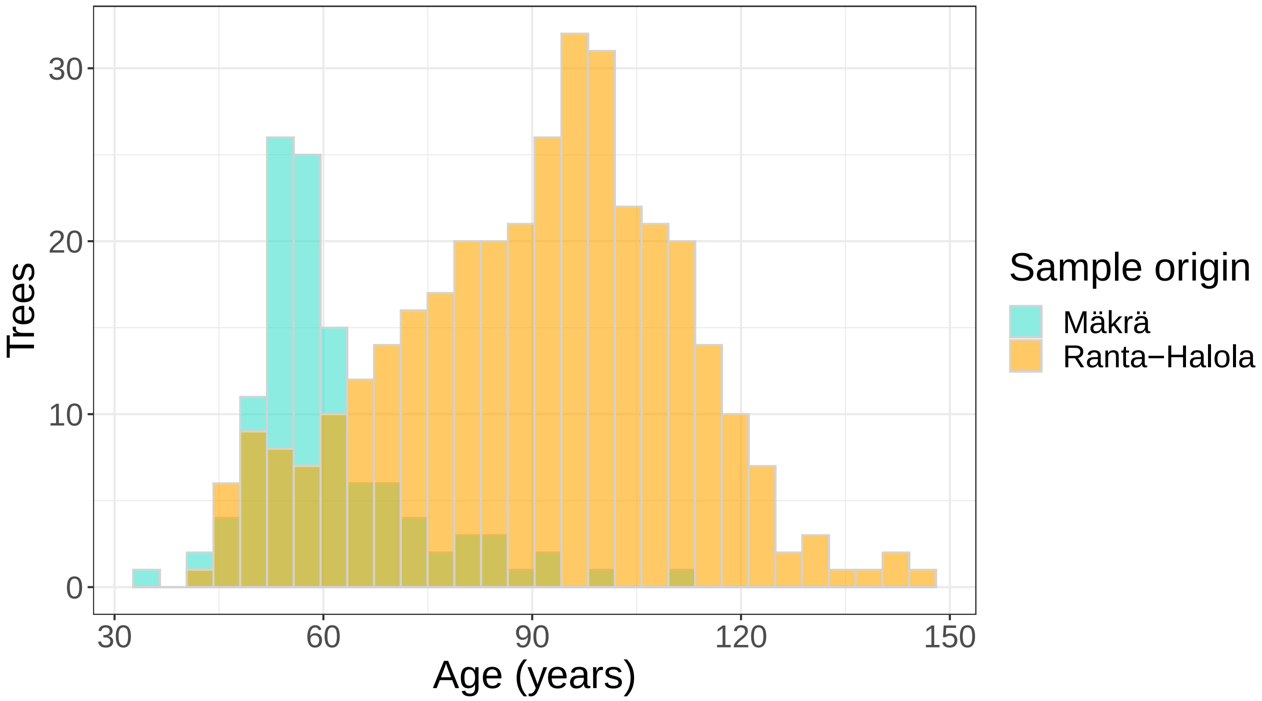


**Figure S1.** The distribution of Scots pine age in Mäkrä (turquoise; *n* = 113) and Ranta-Halola (orange; *n* = 354) sampling sites.

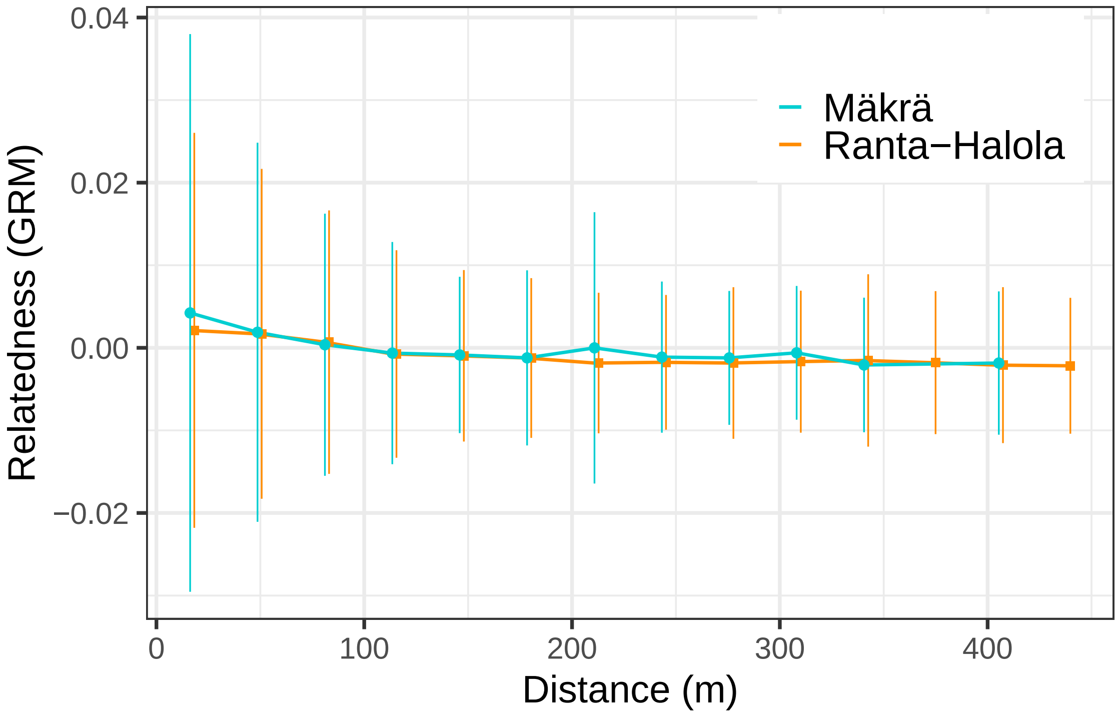


**Figure S2.** Decay of pairwise relatedness with distance in Mäkrä (turquoise) and Ranta-Halola (orange) sampling sites. Mean (circles and squares) and standard deviation (vertical lines) of relatedness is plotted for each distance class; number of pairwise comparisons in each distance class are shown in Table S1. The estimates of Ranta-Halola have been moved two metres forward in the plot to avoid overlap with the estimates of Mäkrä.

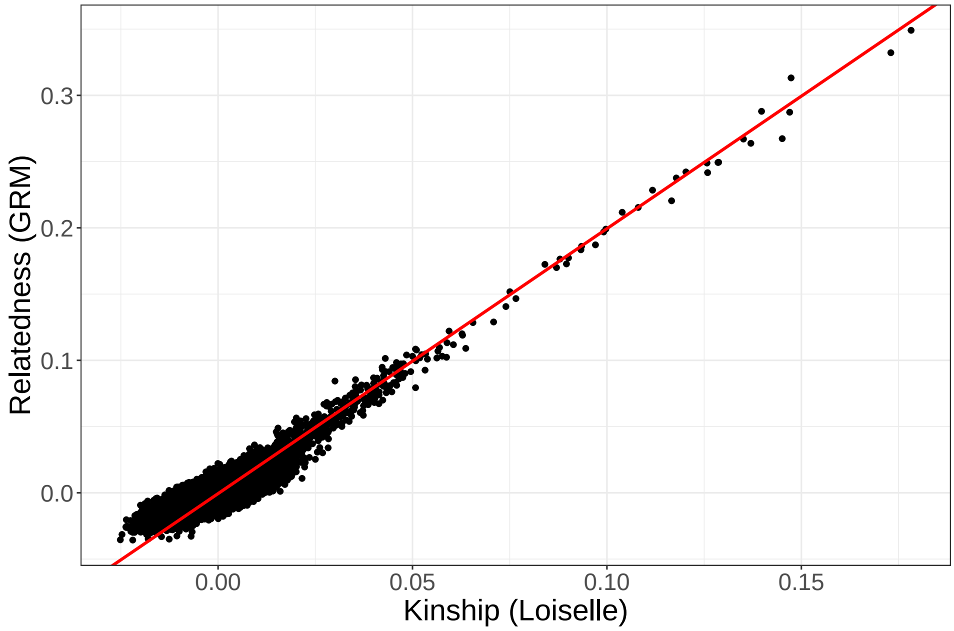


**Figure S3.** Pairwise relatedness (GRM; Yang *et al*. 2011) plotted against pairwise kinship (Loiselle; Loiselle et al. 1995) estimated for 468 Scots pines in the Punkaharju research area. The red line shows the expected relationship of 2:1 for these estimates.

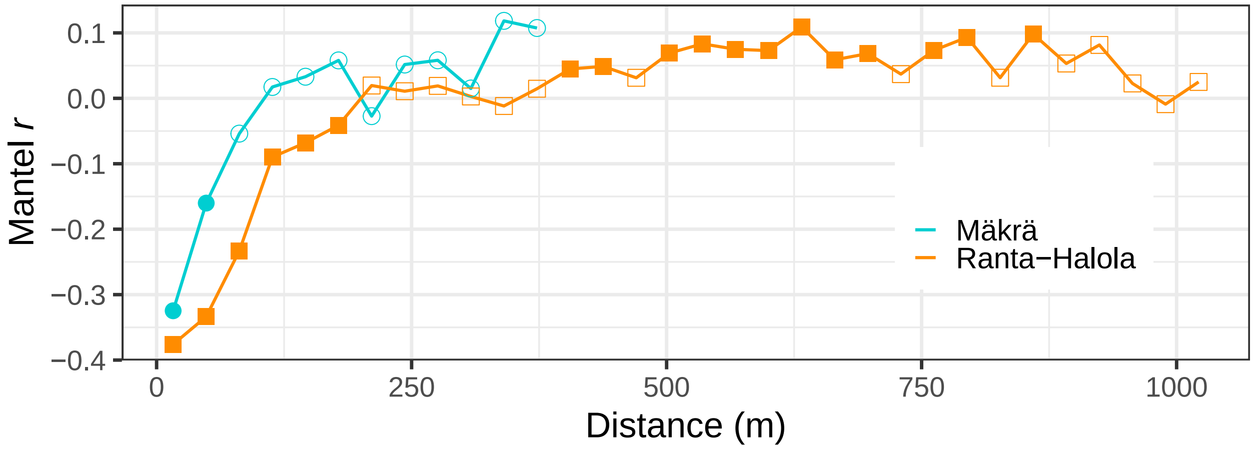


**Figure S4.** Correlation (measured as Mantel *r*) between pairwise relatedness and distance within each distance class estimated as Mantel correlogram in a) Mäkrä (circles, turquoise) and b) Ranta-Halola (squares, orange). Filled shape indicates a *p*-value smaller than 0.05. Ranta-Halola and Mäkrä are divided into equally long distance classes. Two of the longest distance classes from Mäkrä and three from Ranta-Halola have been left out due to including less than 100 pairwise comparisons.

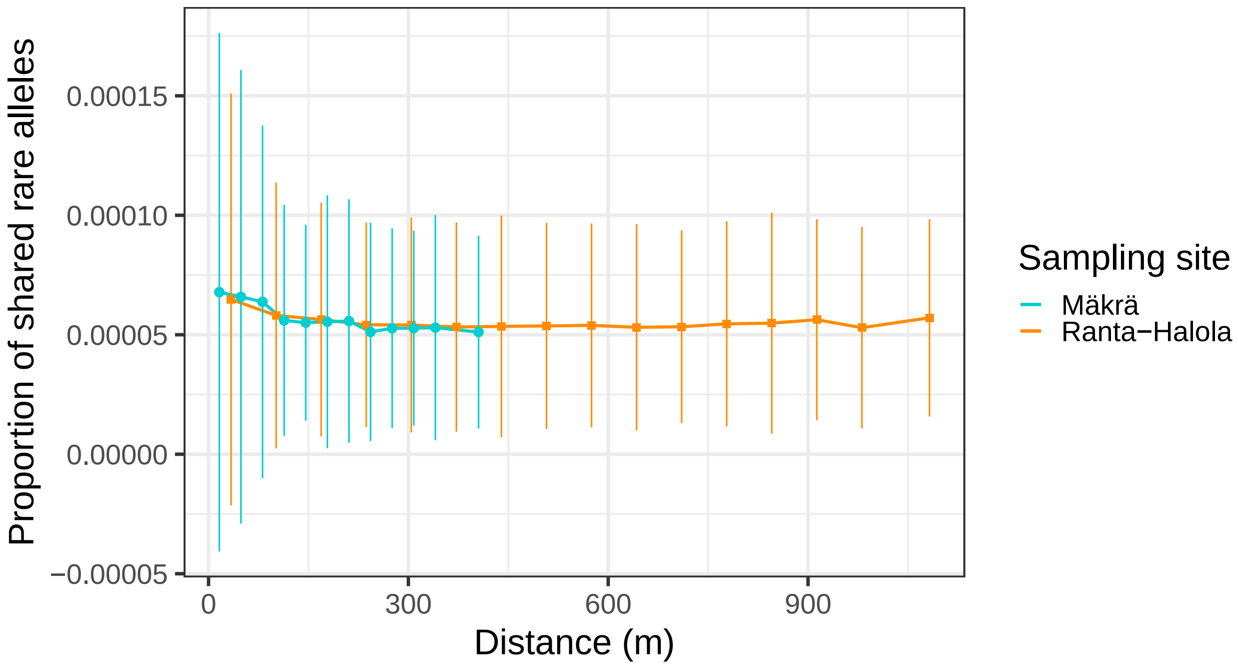


**Figure S5.** Decay of the proportion of shared rare alleles with spatial distance in Mäkrä (turquoise) and Ranta-Halola (orange) sampling sites. Mean (circles and squares) and standard deviation (vertical lines) of relatedness is plotted for each distance class.

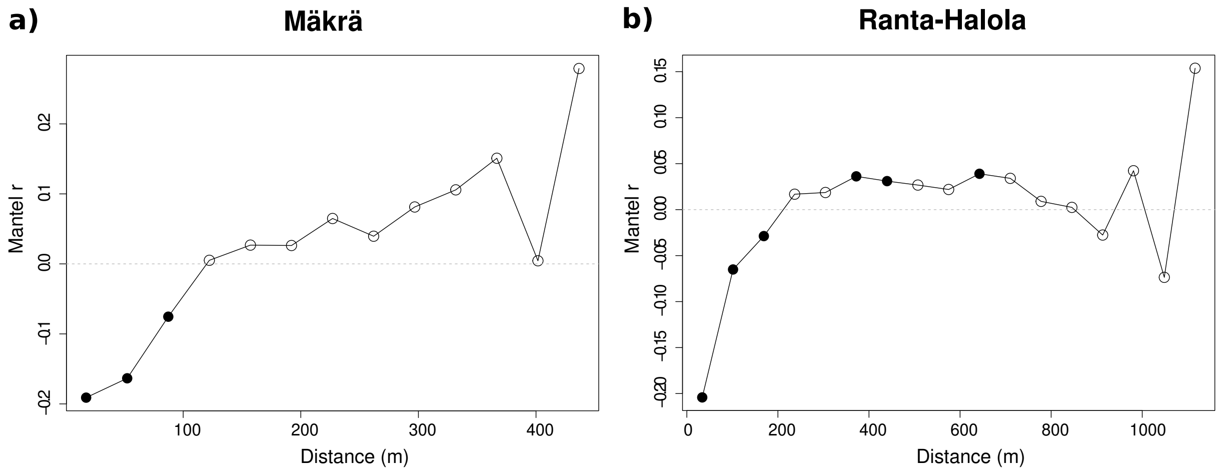


**Figure S6.** Mantel correlogram for rare allele sharing and pairwise distance in a) Mäkrä and b) Ranta-Halola. Filled circles indicate $p$-value smaller than 0.05.

### Supplementary Tables

**Table S1.** The mean and standard deviation (SD) of pairwise relatedness (GRM) in each distance class for Ranta-Halola and Mäkrä sampling sites (Figure 3; Figure S2). The last two and three distance classes for Ranta-Halola (upper panel) and Mäkrä, respectively, are combined so that each class has at least 100 comparisons. The lower panel for Ranta-Halola shows the relatedness values for Ranta-Halola, when distances are classified according to Mäkrä's 14 distance classes.

| | Mean of the distance class (m) | | | | | | | | | | | | | | | |
|---|---|---|---|---|---|---|---|---|---|---|---|---|---|---|---|---|
| | **34** | **101** | **169** | **237** | **304** | **372** | **440** | **507** | **575** | **643** | **710** | **778** | **846** | **913** | **981** | **1082** |
| **Ranta-Halola** | | | | | | | | | | | | | | | | |
| **Mean GRM** | 0.002 | 0.000 | -0.001 | -0.002 | -0.002 | -0.002 | -0.002 | -0.002 | -0.002 | -0.003 | -0.002 | -0.002 | -0.002 | -0.002 | -0.002 | -0.002 |
| **SD GRM** | 0.022 | 0.013 | 0.010 | 0.008 | 0.009 | 0.009 | 0.009 | 0.008 | 0.008 | 0.008 | 0.008 | 0.008 | 0.008 | 0.008 | 0.008 | 0.007 |
| **Pairs** | 2423 | 5832 | 7210 | 7380 | 6938 | 5900 | 5150 | 4419 | 4207 | 3876 | 3401 | 2704 | 1797 | 964 | 437 | 197 |
| | **16** | **49** | **81** | **114** | **146** | **178** | **211** | **243** | **276** | **308** | **341** | **373** | **405** | **438** | | |
| **Mäkrä** | | | | | | | | | | | | | | | | |
| **Mean GRM** | 0.004 | 0.002 | 0.000 | -0.001 | -0.001 | -0.001 | 0.000 | -0.001 | -0.001 | -0.001 | -0.002 | | -0.002 | | | |
| **SD GRM** | 0.034 | 0.023 | 0.016 | 0.013 | 0.009 | 0.011 | 0.016 | 0.009 | 0.008 | 0.008 | 0.008 | | 0.009 | | | |
| **Pairs** | 169 | 498 | 693 | 854 | 858 | 825 | 727 | 601 | 450 | 296 | 168 | | 189 | | | |
| **Ranta-Halola** | | | | | | | | | | | | | | | | |
| **Mean GRM** | 0.002 | 0.002 | 0.001 | -0.001 | -0.001 | -0.001 | -0.002 | -0.002 | -0.002 | -0.002 | -0.002 | -0.002 | -0.002 | -0.002 | | |
| **SD GRM** | 0.024 | 0.020 | 0.016 | 0.013 | 0.010 | 0.010 | 0.009 | 0.008 | 0.009 | 0.009 | 0.010 | 0.009 | 0.009 | 0.008 | | |
| **Pairs** | 571 | 1665 | 2474 | 3000 | 3351 | 3495 | 3538 | 3554 | 3429 | 3334 | 3079 | 2799 | 2700 | 3158 | | |

**Table S2.** Comparison of family relationship classes estimated for 468 Scots pines in the Punkaharju research area using relatedness (GRM; Yang *et al*. 2011) or kinship estimate (Loiselle; Loiselle et al. 1995). Estimates on the darker green background show the same degree of family relationship and ligher green shows one degree difference in the estimated class. Relatedness was estimated using 65 498 SNPs with MAF ≥ 0.05 and kinship using 28 378 SNPs with MAF ≥ 0.20 due to computational reasons.

| | | **Kinship (Loiselle)** | | | | |
|---|---|---|---|---|---|---|
| | | 1st degree | 2nd degree | 3rd degree | 4th degree | unrelated |
| **Relatedness (GRM)** | 1st degree | 0 | 0 | 0 | 0 | 0 |
| | 2nd degree | 1 | 23 | 0 | 0 | 0 |
| | 3rd degree | 0 | 1 | 43 | 6 | 0 |
| | 4th degree | 0 | 0 | 8 | 165 | 24 |
| | unrelated | 0 | 0 | 0 | 32 | 108975 |
